# Let-7b-5p loaded Mesenchymal Stromal Cell Extracellular Vesicles reduce *Pseudomonas*- biofilm formation and inflammation in CF Bronchial Epithelial Cells

**DOI:** 10.1101/2025.05.28.656674

**Authors:** Sharanya Sarkar, Roxanna Barnaby, Amanda Nymon, Lily A. Charpentier, Lily Taub, Matthew J. Wargo, Daniel J. Weiss, Tracey L. Bonfield, Bruce A. Stanton

## Abstract

Cystic Fibrosis (CF) is a multiorgan disease caused by mutations in the *CFTR* gene, leading to chronic pulmonary infections and hyperinflammation. Among pathogens colonizing the CF lung, *Pseudomonas aeruginosa* is predominant, infecting over 50% of adults with CF, and becoming antibiotic-resistant over time. Current therapies for CF, while providing tremendous benefits, fail to eliminate persistent bacterial infections, chronic inflammation, and irreversible lung damage, necessitating novel therapeutic strategies. Our group engineered mesenchymal stromal cell derived extracellular vesicles (MSC EVs) to carry the microRNA let-7b-5p as a dual anti-infective and anti-inflammatory treatment. MSC EVs are low-immunogenicity platforms with innate antimicrobial and immunomodulatory properties, while let-7b-5p reduces biofilm formation and inflammation. In a preclinical CF mice model, we reported that let-7b-5p-loaded MSC EVs reduced *P. aeruginosa* burden, immune cells, and proinflammatory cytokines in the lungs. We hypothesize four complementary mechanisms for the observed *in-vivo* effects of the let-7b-5p loaded MSC EVs: antimicrobial activity, anti-inflammatory properties, inhibition of antibiotic-resistant *P. aeruginosa* biofilm formation in CF airways, and stimulation of anti-inflammatory macrophage behaviors. This study focused on the second and third mechanisms and demonstrates that MSC EVs engineered to contain let-7b-5p effectively blocked the formation of antibiotic-resistant *P. aeruginosa* biofilms on primary human bronchial epithelial cells (pHBECs) while also reducing *P. aeruginosa*-induced inflammation. This approach holds promise for improving outcomes for people with CF. Future work will focus on optimizing delivery strategies and expanding the clinical applicability of MSC EVs to target other CF-associated pathogens.

**NEW AND NOTEWORTHY:** This is the first study demonstrating that let-7b-5p loaded Mesenchymal Stromal Cell Extracellular Vesicles (MSC EVs) block antibiotic-resistant *P. aeruginosa* biofilm formation and reduce inflammation in CF primary human bronchial epithelial cells.

**GRAPHICAL ABSTRACT:** 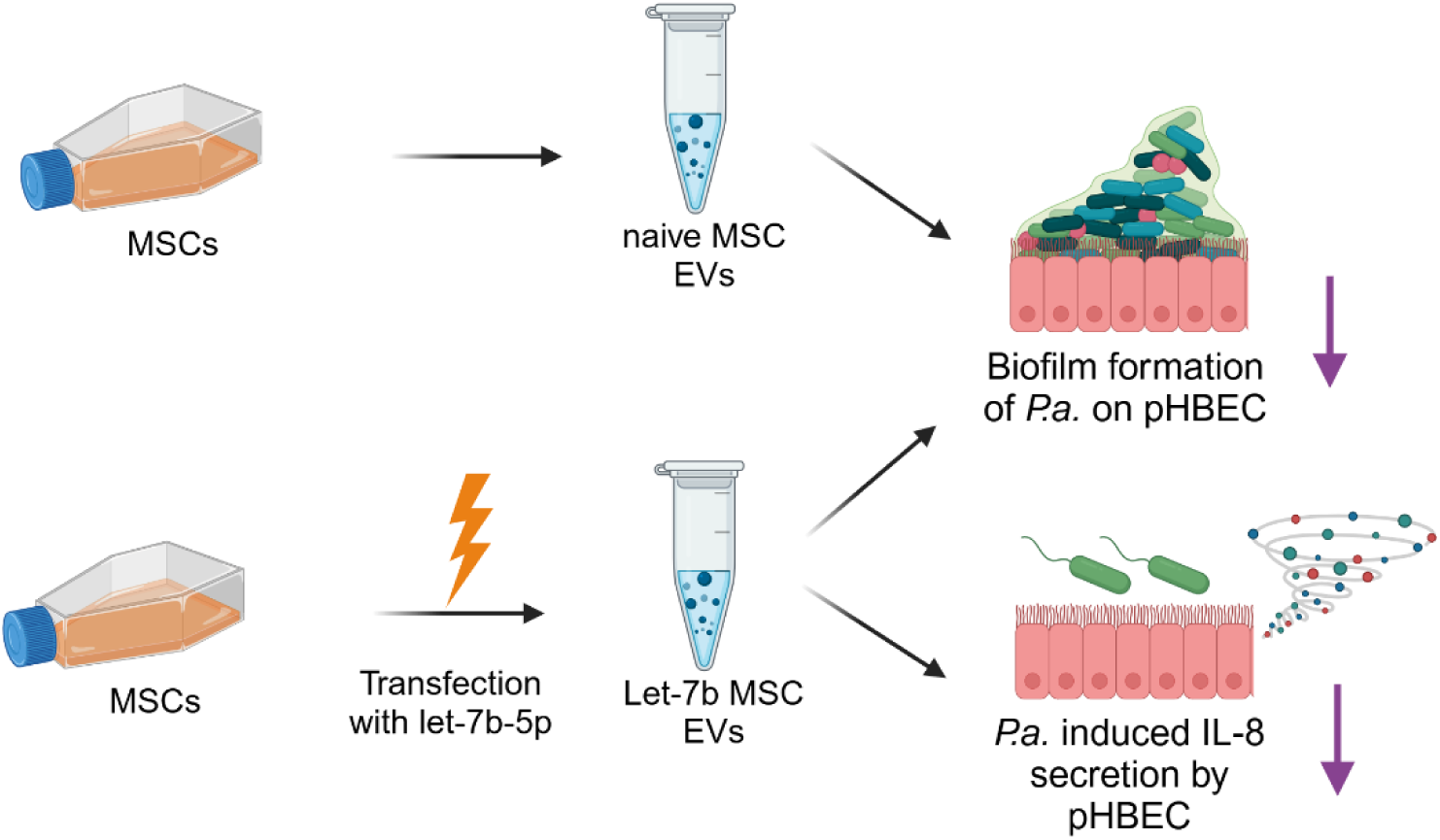

## INTRODUCTION

Cystic Fibrosis (CF), caused by mutations in the cystic fibrosis transmembrane conductance regulator (*CFTR*) gene, is a multi-organ disease that, in the current treatment era, most severely affects the lung. People with CF (pwCF) have impaired mucociliary clearance, resulting in chronic pulmonary infections^1,2^. The CF lung also manifests excessive inflammation, regardless of the status of an infection^3,4^.

Although the CF lung can be colonized by different pathogens including various bacterial, fungal, and viral pathogens, *Pseudomonas aeruginosa (P. aeruginosa)* is the most common pathogen, infecting more than 50% of adult pwCF^5^. Apart from becoming antibiotic-resistant over time, *P. aeruginosa* forms antibiotic resistant aggregates in the airways that function as self-associated biofilms, and are extremely hard to treat once formed^6,7^. Constituents of biofilm structure, such as exopolysaccharides (EPS), are recognized for their ability to sequester reactive oxygen species (ROS), interfere with phagocytic killing, and confer resistance against host antimicrobial peptides^8–10^. Therefore, antibiotic-resistant *P. aeruginosa* biofilms contribute substantially to morbidity and mortality in pwCF^11–14^.

Further, pwCF have inherent defects in their immune cell functions and therefore are unable to clear the persistent infection, exacerbating damage to the lung tissue over time and contributing to progressive lung damage^15^. The CF lung landscape is typically marked by stressed or injured epithelial cells and excessive expression of pro-inflammatory cytokines and alarmins^16–18^.

Although Highly Effective Modulator Therapy (HEMT) such as ETI (Elexacaftor/Tezacaftor/Ivacaftor; Trikafta) improves lung function and reduces hospitalizations in pwCF, it does not fully eliminate bacterial infections or fully ameliorate chronic inflammation^19,20^. HEMTs are available for ∼90% of pwCF, leaving 10% of pwCF few options to reduce infection and inflammation^21^. Furthermore, *P. aeruginosa* reduces the effectiveness of modulators like Orkambi (Lumacaftor/Ivacaftor) that improve CFTR chloride secretion by CF Primary Human Bronchial Epithelial Cells (pHBEC) ^22–24^. Similarly, long-term use of antibiotics and anti-inflammatory drugs in CF poses significant risks^25^. Thus, new treatments are needed to address *P. aeruginosa* infection and inflammation in all pwCF. These approaches may also be applicable to patients with other lung diseases characterized by chronic antibiotic-resistant lung infections such as non-CF bronchiectasis.

In a previous study we demonstrated that the miRNA let-7b-5p, secreted in extracellular vesicles (EVs) by primary wild-type pHBEC, effectively prevents the formation of antibiotic-resistant *P. aeruginosa* biofilms when combined with sub-minimum inhibitory concentrations (MIC) of aztreonam or carbenicillin, antibiotics frequently prescribed for pwCF^26^. This combination also enhances the antibiotic susceptibility of both mucoid and non-mucoid clinical strains of *P. aeruginosa* isolated from pwCF. The underlying mechanism involved a reduction in key proteins associated with biofilm formation and antibiotic efflux pumps in *P. aeruginosa* mediated by let-7b-5p^26^. Additionally, pHBEC-derived EVs increase the sensitivity of *P. aeruginosa* to ciprofloxacin, a commonly used antibiotic in pwCF^27^. We also demonstrated that pHBEC EVs reduce the *P. aeruginosa* load, immune cell infiltration, and pro-inflammatory cytokine levels in CF mouse (gut-corrected F508del/F508del Cftr^tm1Kth^ TG(FABPCFTR)1Jaw/CWR mouse) lungs, even without antibiotics^28^. Furthermore, let-7b-5p is predicted to target other CF-associated pathogens, including Streptococcus and *Burkholderia cenocepacia* and non-tuberculosis mycobacterium (NTM)^29^.

Despite the strong antibacterial effects of pHBEC EVs observed both *in vitro* and *in vivo*, we transitioned to using EVs from Mesenchymal Stromal Cells (MSCs) as an alternative delivery vehicle for let-7b-5p. This shift was prompted by concerns about the potential immunogenicity of pHBEC EVs, which has been noted in some studies^30–33^. In contrast, MSCs, which are precursor cells derived from bone marrow, adipose tissue, or umbilical cord blood, are known for their low immunogenicity ^34^. Importantly, a phase 1 clinical trial involving 15 adult pwCF demonstrated that intravenous infusions of allogeneic human MSCs were safe and well-tolerated^35^. MSCs also have immunomodulatory and antibacterial properties and have been studied for their potential in treating both acute and chronic inflammatory lung diseases^36–38^. Like parent MSCs, MSC EVs have shown good tolerance when administered by aerosol inhalation in healthy volunteers^39^ and have been found to be safe and effective in some populations for treating other diseases^40–42^. MSC EVs have also been shown to be antimicrobial in non-CF contexts. They reduce bacterial load in an *Escherichia coli*-induced pneumonia model^43^ and in *ex-vivo* human lung tissue with severe pneumonia^44^, and were effective in alleviating pulmonary inflammation^45^.

In this report we engineered MSC EVs to contain let-7b-5p as a therapeutic strategy to combat infection and inflammation in CF. We hypothesized that adding let-7b-5p to MSC EVs, which at baseline contain undetectable levels of let-7b-5p^46,47^, would enhance the anti-inflammatory and bactericidal effects of MSC EVs on pHBEC. Our approach combines the pro-resolving, anti-inflammatory properties of MSC EVs with the antibacterial and anti-inflammatory effects of let-7b-5p, thus we predict that adding let-7b-5p will enhance the anti-inflammatory and bactericidal effects of MSC EVs. For example, Ti et al. demonstrated that MSC EVs with elevated levels of let-7b-5p reduce chronic inflammation in a rat wound healing model^48^. Additionally, let-7b-5p is significantly downregulated in bronchial brushings from pwCF compared to non-CF individuals, which may contribute to the hyperinflammatory state observed in CF^49^. Furthermore, let-7b-5p, as a microRNA that targets multiple genes involved in bacterial pathways, may reduce the likelihood of bacterial resistance^29^. Finally, there are currently no registered clinical trials exploring the use of let-7 family members for treating bacterial infections^50^, indicating a notable knowledge gap.

We have demonstrated that both negative control MSC EVs (MSC EVs loaded with a negative control miRNA with no known targets in humans, mice, or *Pseudomonas*) and let-7b-5p-loaded MSC EVs significantly reduced bacterial burden and immune cell infiltration in the lungs of CF mice infected with a clinical mucoid *Pseudomonas* isolate^51^. We hypothesize four mechanisms for the observed *in-vivo* effects of MSC EVs: (i) antimicrobial activity, (ii) anti-inflammatory properties, (iii) inhibition of antibiotic-resistant *P. aeruginosa* biofilm formation, and (iv) promoting CF macrophage phagocytosis to eliminate bacteria and mitigating the inflammatory response of CF macrophages to *P. aeruginosa*. While the first mechanism is well-established, we aimed to explore the second and third mechanisms. In this study, we found that both MSC EVs and let-7b-5p-loaded MSC EVs prevented biofilm formation of *P. aeruginosa* on pHBECs. However, only let-7b-5p-loaded MSC EVs reduced *P. aeruginosa*-induced inflammation in CF-pHBECs. Neither type of MSC EVs increased the intrinsic bacterial killing capacity of CF-pHBEC.

## METHODS

### Culture of Mesenchymal Stromal Cells (MSCs), EV preparation, and miRNA transfection of MSCs

We utilized MSCs from two sources: PACT and ATCC. PACT MSCs were obtained through the NHLBI-funded Production Assistance for Cellular Therapies (PACT) program at the University of Minnesota (Contract: HHSN268201600014I, PI: Dr. David H. McKenna). ATCC Bone Marrow-Derived MSCs were commercially sourced (Cat # PCS-500-012). PACT MSCs were cultured in MEM/EBSS supplemented with 20% FBS and 10% Penicillin-Streptomycin, while ATCC MSCs were maintained following the ATCC’s protocol using their proprietary media with 2% FBS.

EVs were prepared from MSC culture supernatants (passages 2-4, 80% confluency) using the ExoQuick-TC EV isolation kit (Cat # EXOTC50A-1, Systems Biosciences), 48 hours after FBS and antibiotics were removed from the culture media, following the manufacturer’s instructions and as previously described by us^28,51^. MSC supernatants were centrifuged and concentrated using 30 kDa filters. The concentrate was combined with ExoQuick-TC polymer, incubated overnight at 4°C, centrifuged, and resuspended in 150 μL PBS. Unconditioned media, not exposed to cells, underwent the same preparation procedure and was used as a process control (PC) to account for potential contaminants in the media and those that may be introduced during EV preparation.

After determining that ATCC MSCs were both purer and more anti-inflammatory compared to PACT MSC EVs, we selected ATCC cells for further experiments. MSCs were transfected with either a negative control (NC) miRNA, that has no known target genes in mice, humans, or *P. aeruginosa*, or let-7b-5p using Lipofectamine RNAiMAX Transfection Reagent (Invitrogen, Waltham, MA), using a protocol previously described by us^51^. Untreated MSCs served as a second set of controls. Media was collected one day post-transfection, and EVs were isolated using ExoQuick-TC, The presence of let-7b-5p in EVs was confirmed via qRT-PCR, as described in our prior work^51^.

### Characterization of PACT and ATCC MSC EVs

MSC EVs secreted by PACT and ATCC cells were analyzed using the Exosome Analytical Services from RoosterBio Inc. (Frederick, MD) with their proprietary assays. Sample identities were blinded to avoid potential bias. Particle concentration and size were measured using Nanosight NS300 with a 532 nm laser, Camera Level 14, Detection Threshold 7, variable focus, and Screen Gain 10. Three 30-second videos were analyzed with the instrument’s software. Protein concentration was determined using a BCA assay, while total lipids were measured with MEMGlow 488, a fluorescent dye that binds to EV lipid bilayers. A standard curve in the sample matrix quantified lipid content. RNA was isolated using the RNeasy Mini Kit, and miRNA concentration was assessed with the 2100 Bioanalyzer and Small RNA Kit. EV purity was determined by staining samples with MEMGlow 488 and comparing stained and unstained particle concentrations using fluorescence detection and standard Nanoparticle Tracking Analysis (NTA) to calculate the percentage of lipid-positive particles.

### Culture of Human Bronchial Epithelial Cells (HBEC)

Primary HBECs (pHBEC) from explanted lungs of three deidentified CF donors homozygous for the ΔF508 mutation undergoing lung transplantation were obtained from Dr. Scott Randell at the University of North Carolina at Chapel Hill (Chapel Hill, NC). The cells were cultured in air-liquid interface (ALI) conditions, following established protocols, and verified to be free of mycoplasma contamination^52,53^. RNA sequencing confirmed the cells were primary human airway epithelial cells. Of the three donors, two were females and one was male, with ages ranging from 14 - 27 years. All donors were nonsmokers. The sample size (n) refers to biological replicates. In brief, pHBEC were maintained in BronchiaLife basal medium (Lifeline Cell Technology, Frederick, MD, Cat. No. LM-0007) at 37°C with 5% CO₂. The medium was supplemented with the BronchiaLife B/T LifeFactors Kit (Lifeline Cell Technology, Cat. No. LS-1047) and antibiotics (penicillin, 10,000 U/mL; streptomycin, 10,000 μg/mL; Sigma-Aldrich, Cat. No. P4333). Cells were used for experiments between passages 4 and 9, and the data was not dependent on the passage number.

### IRB statement

The Dartmouth Committee for the Protection of Human Subjects has concluded that this study’s use of MSCs and pHBEC does not qualify as human subject research since the cells were derived from discarded tissue and lack any patient identifiers. All tissues utilized for cell isolation were collected with informed consent, ensuring the donor’s personal information remains confidential.

### P. aeruginosa cultures

*P. aeruginosa* strain PA14 was retrieved from frozen glycerol stock and cultured in Luria broth (LB, Thermo Fisher Scientific, Waltham, MA) at 37°C with shaking at 225 rpm for 16 hours. Following this initial incubation, 5 µL of the overnight culture was transferred into 5 mL of fresh LB media and incubated under the same conditions for an additional 16 hours to ensure robust bacterial growth. After the second overnight culture, the bacteria were washed twice with PBS (Gibco) to remove residual media. The optical density at 600 nm (OD₆₀₀) was measured, and the absorbance values were applied to a standard curve (OD₆₀₀ vs. CFU/mL) established in our lab to calculate the required volume for achieving the desired inoculum (CFU/mL) in subsequent experiments.

### Treatment of CF-pHBEC with *P. aeruginosa* and MSC EVs

CF-pHBECs (5 × 10⁵) derived from 3 CF donors (ΔF508/ΔF508) were individually seeded on 24-mm permeable supports (Corning, Catalog #3407) pre-coated with type IV collagen (Sigma-Aldrich, Catalog #C-7521). The cells were cultured in air-liquid interface (ALI) media (University of North Carolina, Chapel Hill, NC) at 37°C for 3 - 4 weeks to establish polarized monolayers^54^. *P. aeruginosa* PA14 (3 × 10⁸/mL) was introduced to the apical side of the monolayers for 1 hour. Subsequently, either process control (PC), NC MSC EVs (2 × 10⁹/mL), or let-7b MSC EVs (2 × 10⁹/mL) were added for an additional 5 hours, completing a 6-hour treatment period. We chose this time frame due to the established cytotoxicity of *P. aeruginosa* towards pHBECs after 6 hours^26,55^. One monolayer of each donor of CF-pHBECs served as a second control, receiving neither *P. aeruginosa* nor EVs, while another monolayer was treated only with *P. aeruginosa* with no EVs. For each CF donor/pass, four 24-mm supports were prepared. The first plate was used to measure cytokine levels by ELISA and extract RNA for qRT-PCR and RNA sequencing. The second plate was used to assess cytotoxicity by measuring LDH in basolateral fluid and cell lysates. The third plate was used to measure bacterial burden by harvesting apical and basolateral fluids along with cell lysates. The fourth plate also assessed bacterial burden but included a pre-treatment with ETI.

For the ETI pre-treatment, Ivacaftor (10 nM; Selleck Chemicals, Catalog #S1144), elexacaftor (3 µM; Selleck Chemicals, Catalog #S8851), and tezacaftor (3 µM; Selleck Chemicals, Catalog #S7059) or DMSO as a vehicle control (Fisher Scientific, Catalog #D128-500) were added to the basolateral media 48 hours before the experiment. ETI was applied for 24 hours, after which the media was replaced with fresh media containing ETI for another 24 hours, totaling 48 hours of ETI exposure prior to starting the infection and treatment experiments.

### Cytotoxicity

Cytotoxicity of treatments was assessed using the CytoTox 96® Non-Radioactive LDH Cytotoxicity Assay (Promega, catalog #G1780) according to the manufacturer’s instructions. Briefly, 50 µL of basolateral fluid from each treatment well was transferred to a 96-well plate, along with 50 µL of unconditioned media as a blank at the 6-hour treatment endpoint. To lyse CF-pHBEC, 10X lysis solution was added at a 1:10 ratio and incubated for 45 minutes at 37°C. Subsequently, 50 µL of the lysates was added to the 96-well plate. CytoTox reagent (50 µL) was added to each well, and the plate was incubated in the dark for 30 minutes, followed by the addition of 50 µL STOP solution to terminate the reaction. Absorbance was measured at 490 nm within 1 hour. Background values from the blank were subtracted, and percent cytotoxicity was calculated by dividing the absorbance of the basolateral fluid to that of the cell lysate.

### Biotic biofilm formation assay

The biotic biofilm formation assay was performed following our previously published protocol^26^. In summary, CF-pHBECs from three donors were cultured as confluent monolayers on glass coverslips. Fluorescent *P. aeruginosa* PA14 (PA14-mKO2: 3 × 10⁶/mL), a strain developed by Dr. Carey Nadell and Swetha Kasetty at Dartmouth College, was pre-exposed to process control (PC), MSC EVs or let-7b-5p MSC EVs for 18 hours. The 18-hour incubation allowed sufficient time for MSC EVs to deliver their cargo to *P. aeruginosa*, and suppress the abundance of proteins critical for biofilm development^26^. The EV-to-bacterium ratio was 1,000:1, a concentration ratio significantly lower than the EV levels observed in CF-pHBEC culture supernatants and human bronchoalveolar lavage fluid (10¹¹/mL)^56,57^.

After this pre-exposure, *P. aeruginosa* were introduced to the apical side of the pHBEC monolayers. The bacteria were allowed 1 hour to adhere to the pHBEC monolayers and imaged over 5 hours. Both the 18-hour pre-exposure and the 6-hour co-culture included a low concentration of aztreonam (0.1 µg/mL, one-half the minimal inhibitory concentration^26^), which alone does not impact planktonic growth or biofilm formation *in P. aeruginosa*. Previously, we demonstrated that aztreonam and EVs secreted by pHBEC have an additive effect to prevent biofilm formation^26^. Imaging was performed as previously described^26^. Z-stack images from 4-5 randomly selected areas of each pHBEC monolayer were captured at 60X magnification using the Keyence BZX800 analyzer software. Three-dimensional volume renderings of PA14-mKO2 fluorescence in the TRITC channel were generated using the software’s 3D Cell Count feature. Confocal Z-stack images were analyzed to calculate total biofilm volumes in the 4-5 random areas per sample by two different investigators, one blinded to the treatment, adhering to the manufacturer’s guidelines for Keyence software. Biofilm CFU/mL were assessed by scraping the pHBECs, and plating the bacteria on LB agar, as described in the section below.

### Assessing total bacterial burden on CF-pHBECs

At the 6-hour time point, 500 µL of apical supernatant and 500 µL of basolateral supernatant were collected from the CF-pHBEC cultures. The cells were gently washed with PBS++ (Gibco) to remove any residual media and debris. Subsequently, 500 µL of 0.1% Triton X-100 (Bio-Rad) was added to the cells, and the plate was placed on a shaker for 15 minutes to facilitate cell lysis. The lysates were then carefully scraped from the filters, transferred to fresh tubes, and mixed thoroughly using a vortex. To quantify bacterial burden, dilution plating was performed for all experimental conditions. 10 µL of appropriate dilutions of the samples were spread onto LB agar plates, which were then inverted and incubated at 37°C for 16 hours. After incubation, colonies were counted, and the counts were adjusted using the respective dilution factors. The bacterial counts from the lysate and planktonic fractions (apical and basolateral supernatants) were combined for each sample to calculate the total bacterial burden, expressed as CFU/mL.

### Measurement of IL-6 and IL-8 secretion by ELISA

The secretion of IL-6 and IL-8 by pHBECs at 6 hours was quantified using the Human IL-6 DuoSet ELISA (Catalog #DY206, R&D Systems, Minneapolis, MN) and Human IL-8/CXCL8 DuoSet ELISA (Catalog #DY208, R&D Systems, Minneapolis, MN), respectively, according to the manufacturer’s protocols. Triplicate wells were analyzed for each sample, as recommended by R&D Systems. Concentrations of IL-6 and IL-8 were determined based on standard curves generated from known concentrations of the analytes. To calculate total secretion, the concentrations from the apical and basolateral compartments of each sample were combined.

### qRT-PCR to assess IL-6 and IL-8 mRNA in CF-pHBECs

RNA was isolated from HBECs using the miRNeasy Kit (Qiagen, Germantown, MD, Catalog #217004) after the 6-hour timepoint. cDNA synthesis was performed with 1 µg of RNA as input, using SuperScript IV (Invitrogen, Grand Island, NY, Catalog #18090050). qRT-PCR was carried out with template cDNA at a concentration of 10 ng/µL, TaqMan Master Mix (Invitrogen, Waltham, MA, Catalog #4304437), and TaqMan gene expression assays for IL-8 (Catalog #Hs00174103_m1, Thermo Fisher Scientific) and IL-6 (Catalog #Hs00985639_m1, Thermo Fisher Scientific), following the manufacturers’ protocols. UBC (Catalog #Hs00824723_m1, Thermo Fisher Scientific) was used as the reference gene, after examining several reference genes to choose one that did not respond to treatment. Cycle threshold (Ct) values were measured in triplicate for each gene and condition on the same PCR plate, and the averages were used for analysis. Fold changes of gene expression were calculated using the 2^-ΔΔCt^ method. Briefly, the Ct value of the target gene (IL-6 or IL-8) was normalized to that of the reference gene (UBC) for both treated and untreated samples (ΔCt = Ct_target - Ct_reference). The ΔCt of the untreated sample (control CF-pHBEC) was then subtracted from the ΔCt of the treated sample (CF-pHBEC treated with *P. aeruginosa*, or PC, or NC MSC EVs, or let-7b MSC EVs) to calculate ΔΔCt (ΔΔCt = ΔCt_treated - ΔCt_untreated). Relative expression was then determined using the formula 2^-ΔΔCt^.

### RNA seq and downstream analyses

RNA for sequencing was quantified using Qubit, and RNA integrity was assessed with a Fragment Analyzer (Agilent). A total of 100 ng of RNA was used as input for library preparation following the QuantSeq FWD workflow (Lexogen) according to the manufacturer’s protocol. Prepared libraries were pooled and sequenced on a NextSeq2000 platform, aiming at 10 million single-end 100 bp reads per sample. Paired-end reads were processed by trimming with Cutadapt v4.0^58^ followed by alignment to the GRCh38 human genome reference using Hisat2 v2.2.1 (RRID:SCR_015530)^59^. Gene expression levels were quantified with FeatureCounts v2.0.1^60^, using Human Ensembl v97 as the reference annotation. Alignment and quantification metrics were assessed with Samtools v1.15.1 (RRID:SCR_001105)^61^ and Picard v2.27.1^62^. Filtering, normalization, and differential gene expression were performed using edgeR v4.2.0^63^ in R v4.4.0^64^. Using the filterByExpr function, transcripts with fewer than 10 counts in each sample were filtered out, retaining 14,165 genes for analysis. Library sizes were adjusted by normalization factors calculated with calcNormFactors. Differential gene expression was determined using gene-wise negative binomial generalized linear models. Figures were created using ggplot2 v3.5.1^65^. Sequencing data has been deposited to NCBI GEO (accession #GSE295607). KEGG pathway analysis was performed using the gseKEGG function of clusterProfiler v4.12.0^66^, with a minimum geneSet size of 120. Significantly differentially expressed genes used for this analysis were defined as having an absolute log_2_ fold change > 1 and an unadjusted P value < 0.05. IntaRNA^67,68^ was used to calculate interaction energies between miRNAs and human RNAs.

### Statistics

Data was analyzed using the R statistical platform (v4.4.0)^64^. Most figures were created using GraphPad Prism (v10.3.1, San Diego, CA), with ggplot2 used for RNA-seq figures (as mentioned above). The figure legends provide details of the respective statistical analyses and include precise p-values. In brief, differential gene expression for RNA-seq data was determined using gene-wise negative binomial generalized linear models. Other analyses were performed using linear mixed-effects models because it allows for the modeling of donor as a random effect and accounts for intrinsic donor variability. For biofilm volumes, interaction analyses between time and treatment were also included into the linear mixed-effects model. The lmerTest^69^ and nlme^70^ packages were used to fit the models and calculate p-values. The sample size (n) represents the number of donors.

### BioRender

The graphical abstract was created using the Premium version of BioRender with the following publication license: Sarkar, S. (2025) https://BioRender.com/i41b371

## RESULTS

### Preparation and characterization of MSC EVs

In our previous study, we showed that MSC EVs reduced bacterial burden in CF mice using EVs derived from ATCC human bone marrow MSCs (Cat # PCS-500-012)^51^. However, given that the purity and potency of MSC EVs can vary depending on their source, we first sought to identify the most effective MSC EVs^71^. Additionally, we aimed to align our study with the International Society for Extracellular Vesicles (ISEV) initiative to strengthen the rigor of EV research^72^. To this end, we conducted a comparative analysis of ATCC MSC EVs and MSC EVs derived from the NIH PACT program at the University of Minnesota (PACT EVs). Both ATCC and PACT MSCs are bone marrow-derived, and the EVs from these two sources were matched by cell passage (i.e. same cell passage) and isolated using the same protocol (ExoQuick-TC) to control for potential confounding factors and ensure a fair comparison of their purity and potency. EVs from ATCC and the PACT were characterized according to ISEV recommendations, assessing: (i) particle concentration, (ii) protein concentration, (iii) lipid concentration, (iv) miRNA content, (v) EV purity, and (vi) size distribution **(Table 1)**. It is noteworthy that ATCC EVs had a higher purity (∼46%) relative to PACT EVs (∼20%), measured by comparing the number of particles labeled with a lipid dye compared to total particle count as determined by Nanoparticle Tracking Analysis (NTA)^72^. The sizes of both types of EVs fell within the range specified by MISEV^72^. Visualization of EVs via Transmission Electron Microscopy (TEM) was performed as described and published in our previous study^51^. In initial screening studies utilizing CF-pHBEC, we observed that ATCC EVs significantly reduced IL-6, as compared to control/untreated cells (p-value = 0.0020), while PACT EVs did not **(Figure 1A)**. Moreover, ATCC EVs (p-value = 0.0004) were better than PACT EVs (p-value = 0.0086) in reducing IL-8, as compared to control **(Figure 1B)**.

**Figure 1:**
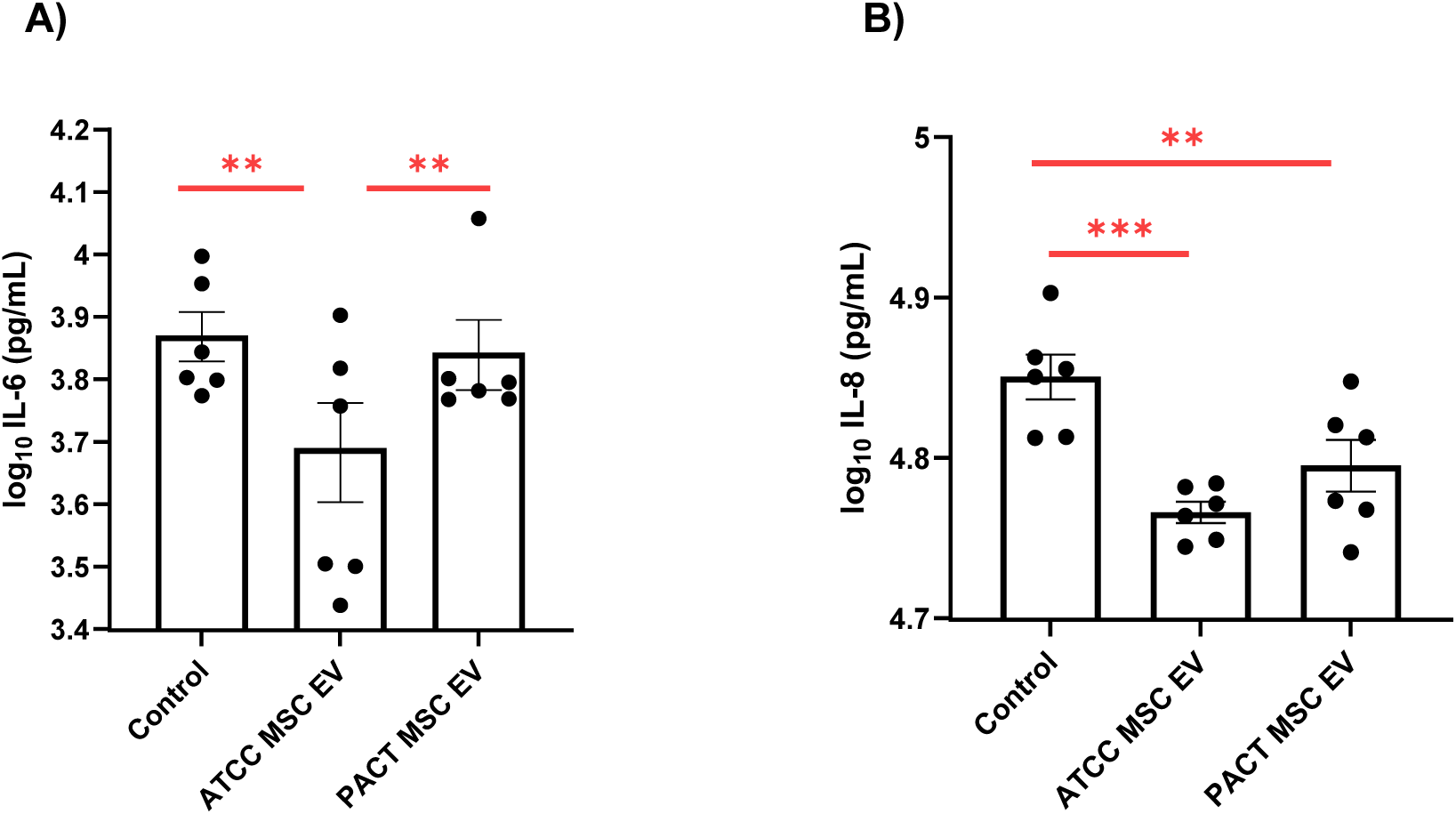
A) ATCC MSC EVs were significantly better at reducing IL-6 secretion by CF-pHBEC compared to PACT MSC EVs (p = 0.0020). B) ATCC MSC EVs (p = 0.0004) were more potent than PACT MSC EVs (p = 0.0086) in reducing IL-8 secretion by CF-pHBEC. CF-pHBEC were exposed to MSC EVs for 6 hours and then apical and basolateral supernatants from CF-pHBEC were collected and analyzed by ELISA. The experiments included CF-pHBEC from two donors (ΔF508/ΔF508), each with 3 replicates (yielding 6 data points per treatment group). Linear mixed-effects models with the donor as a random effect were used to test the statistical significance between the control and EV groups. **p < 0.01; ***p < 0.001; mean ± SEM depicted.

**Table 1:** Characterization of ATCC and PACT MSC EVs.

| Sample | Particle concentration (particles/mL) | Total protein ± SD (μg/mL) | Total lipid ± SD (μg/mL) | Total miRNA ± SD (pg/μL) | EV purity (%) | Mean particle size ± SD (nm) |
| --- | --- | --- | --- | --- | --- | --- |
| ATCC | $6.8 \times 10^9$ | $24.7 \pm 1.2$ | $14.6 \pm 0.4$ | $147.9 \pm 32.5$ | 46.4 | $151.1 \pm 64.4$ |
| PACT | $8.3 \times 10^9$ | $85.5 \pm 0.3$ | $32.7 \pm 3.9$ | $130.4 \pm 92.9$ | 20.5 | $133.6 \pm 52.4$ |

### *P. aeruginosa* is not cytotoxic to CF-pHBEC

Due to their superior purity and potency, we employed ATCC MSC EVs in all future studies in this report. ATCC MSCs were transfected with either a negative control (NC) miRNA (a miRNA with no known targets in humans, mice, or *Pseudomonas*) or let-7b-5p miRNA. Unconditioned MSC media processed through the EV preparation protocol served as the process control (PC) to account for any possible effects on CF-pHBEC due to the MSC media or EV preparation procedure. qRT-PCR was used to confirm let-7b loading in MSC EVs^51^. Next, we assessed whether *P. aeruginosa* was cytotoxic to CF-pHBEC at a 6-hour time point, either alone or in combination with PC, NC MSC EVs, or let-7b MSC EVs. LDH assays revealed that none of the treatments altered LDH release compared to control (untreated) cells **(Figure 2)**. While cytotoxicity varied across different donors, reflecting inherent donor-specific differences, typical in pwCF, the LDH values were consistent across all treatments within each donor.

**Figure 2:**
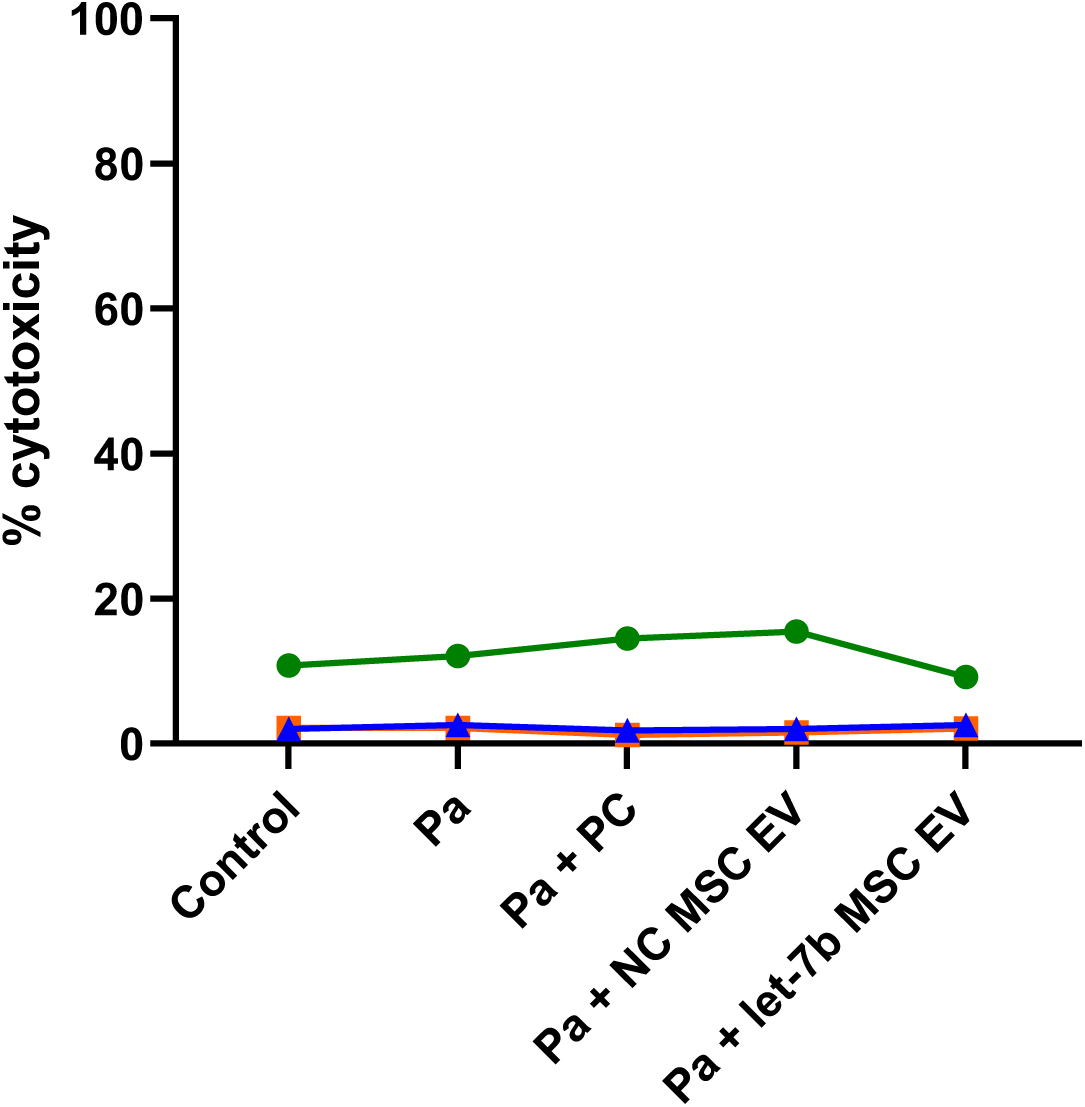
*P. aeruginosa* (Pa) alone or in combination with PC, NC MSC EVs, or let-7b MSC EVs were not cytotoxic as compared to untreated CF-pHBEC 6 hours after exposure. Each colored line represents one CF donor across all treatment conditions. Linear mixed-effects models with donor as random effect were used to test for statistical significance between untreated and the treatment groups. n = 3 CF donors (ΔF508/ΔF508). PC (process control) = unconditioned media passed though EV preparation process, NC = negative control miRNA.

### MSC EVs Inhibit Biofilm Formation by *P. aeruginosa* on pHBEC

To test the hypothesis that MSC EVs inhibit the ability of *P. aeruginosa* to form biofilms, experiments were conducted using fluorescent *Pseudomonas* PA14-mKO2 added to the apical side of pHBEC^26^. As described in detail in Methods, *P. aeruginosa* was pre-exposed to either PC, MSC EVs, or let-7b MSC EVs for 18 hours. This pre-incubation period allowed sufficient time for the MSC EVs to deliver their cargo to the bacteria and reduce the abundance of proteins essential for biofilm formation as shown previously for EVs secreted by pHBEC^26^. Following the pre-incubation, the bacteria were added to pHBEC monolayers, allowed 1h to attach to the cells, and were then imaged over the next 5h period by fluorescence microscopy, since longer exposure of *P. aeruginosa* is cytotoxic^26,55^. Imaging revealed that attachment of *P. aeruginosa* (1 hour) was similar in both treatment groups **(Figures 3A - 3C)**. PC pre-treated *P. aeruginosa* developed robust biofilms in 6 hours (p-value = 5.02 × 10^-7^) **(Figures 3A - 3B)**, whereas MSC EVs blocked the formation of biofilms **(Figures 3C - 3D)**. Let-7b-5p-loaded MSC EVs also completely inhibited biofilm formation **(Figure 3E)**. Taking an orthogonal approach to measure biofilm formation by microscopy, we removed planktonic bacteria and scraped the biofilm *P. aeruginosa* at the end of the experiments and plated to count colony forming units (CFU). The initial inoculum of *P. aeruginosa* added to the cells at the beginning of the assay was 2.00 × 10^6^ CFU/mL. We observed that *P. aeruginosa* pre-treated with PC increased biofilm *P. aeruginosa* to 1.08 × 10⁷ CFU/mL (a 440% increase from the initial inoculum) at the end of the 6-hour experiment, consistent with the fluorescent images of biofilms. By contrast, both MSC EV and let-7b MSC EV dramatically reduced the increase in CFUs. *P. aeruginosa* biofilm CFUs were 2.84 × 10⁶ CFU/mL in NC MSC EV exposed *P. aeruginosa* (a 42% increase from the initial inoculum) and were 2.77 × 10⁶ CFU/mL in let-7b-5p-MSC EV exposed *P. aeruginosa* (a 38.5% increase from the initial inoculum) respectively, considerably less than the 440% increase in *P. aeruginosa* biofilm CFUs (p-value = 0.023 and 0.005, respectively) **(Figure 3F)**. Moreover the CFUs at the end of the MSC EV exposure were statistically similar to the input of 2.00 × 10^6^ CFU/mL.

**Figure 3:**
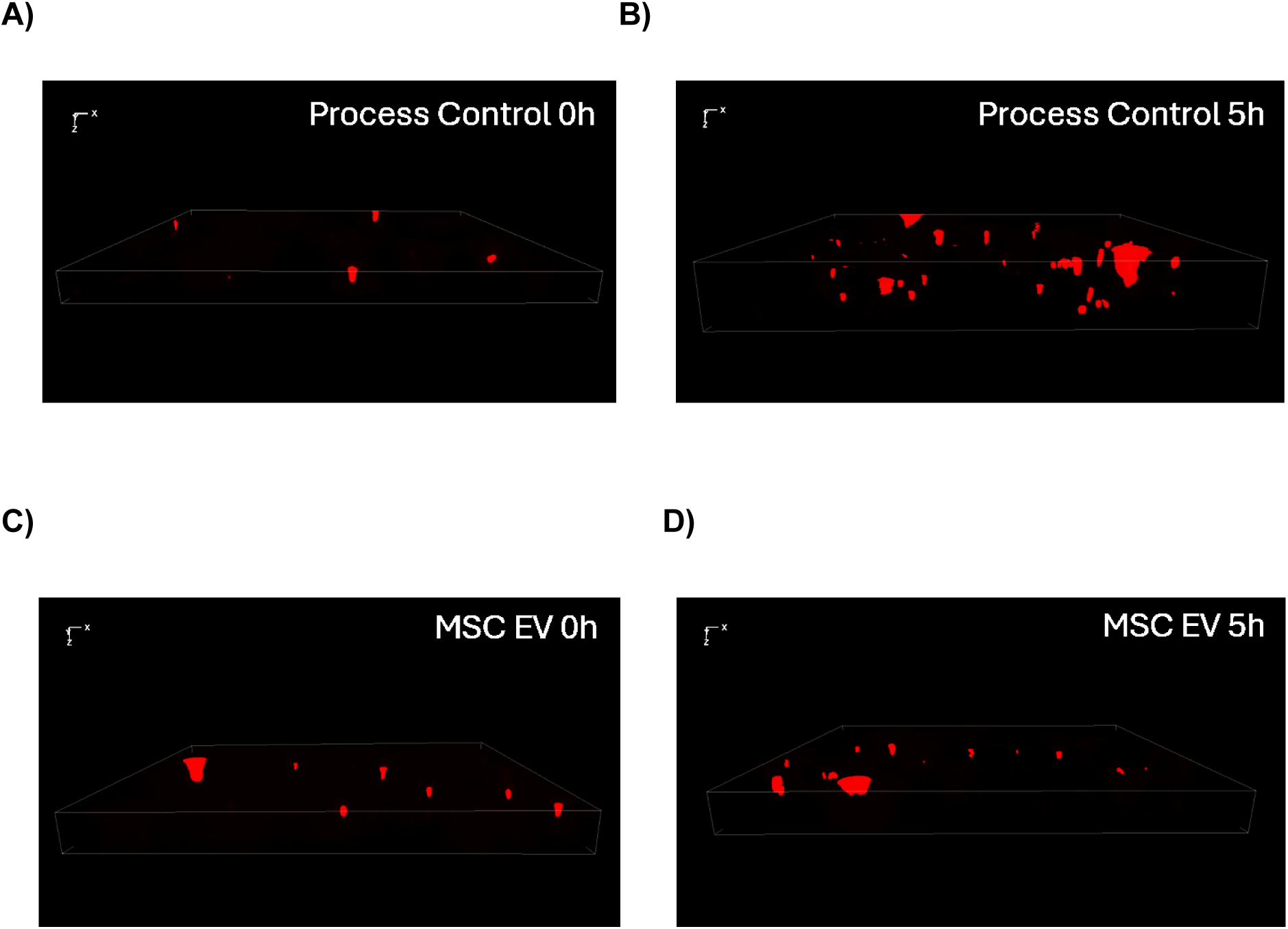
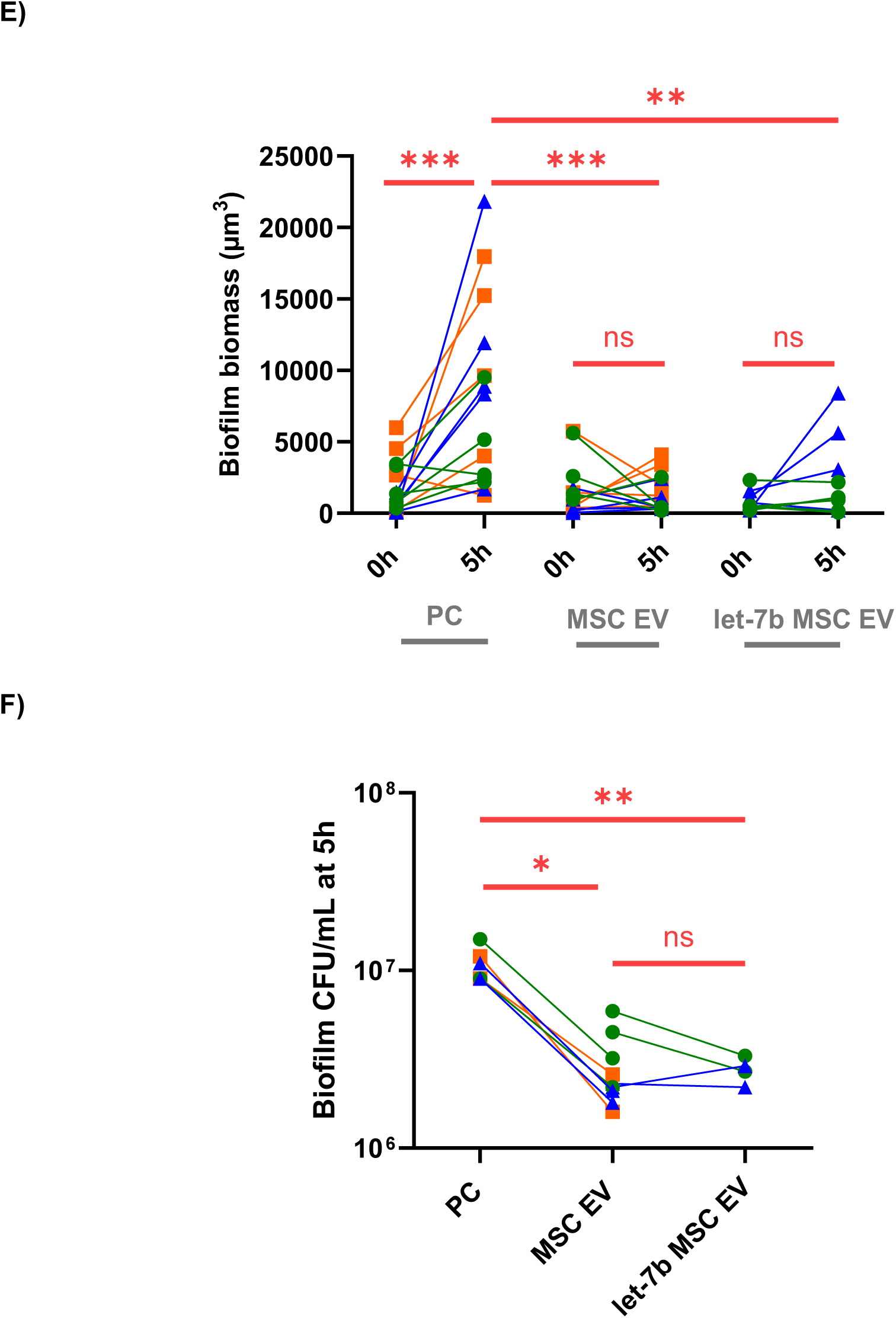
Representative images of maximum intensity projections from z-stacks of biotic biofilms of *P. aeruginosa* pre-treated with PC at (A) 0h and at B) 5h after the 1 hour of exposure to enable the bacteria to attach to pHBEC. Panels C and D are images of *P. aeruginosa* pre-treated with MSC EVs at 0h (C) and at 5h (D) on pHBEC monolayers. (E) Summary of data of biofilm volumes of *P. aeruginosa* exposed to PC, MSC EVs or let-7b-5p MSC EVs on pHBEC. Volume renderings of 3D Z-stacks from 1-5 random areas of each monolayer were generated using Keyence software following the manufacturer’s instructions. PC pre-treated *P. aeruginosa* significantly increased biofilm volume over 5h (p =5.02 × 10^-7^), whereas there was no significant increase in biofilm volumes of *P. aeruginosa* pre-treated with MSC EVs or let-7b MSC EVs. MSC EV pre-treated *P. aeruginosa* and let-7b MSC EV pre-treated *P. aeruginosa* have significantly lower (p = 0.000163 and 0.00762, respectively) biofilm volumes compared to PC pre-treated *P. aeruginosa* at 5h. There was no difference in the attachment of *P. aeruginosa* to pHBEC at 0 hours in all three treatments. Each data point represents the *P. aeruginosa* biofilm volume on a monolayer from an individual pHBEC donor, with colored lines connecting measurements from the same donor at 0 and 5h. Each colored line represents a unique pHBEC donor, and lines of the same color are technical replicates (different imaged areas) of the same donor. (F) MSC EV pre-treated *P. aeruginosa* and let-7b MSC EV pre-treated *P. aeruginosa* have significantly lower (p = 0.023 and 0.005, respectively) biofilm CFU/mL compared to PC pre-treated *P. aeruginosa* at 5h. The lines connect CFUs formed on monolayers from each donor across the different pre-treatment conditions. Colored lines and symbols represent replicates of the same donor. Linear mixed-effects models with donor as random effect were used to test statistical significance, and for the biofilm volumes, interaction analyses between time and treatment (to determine if the treatment effect was modified by time) were also included in the model; n = 3 pHBEC donors; *p < 0.05, **p < 0.01; ***p < 0.001; PC (process control) = unconditioned media passed through the EV preparation process.

### Let-7b-5p MSC EVs reduce *Pseudomonas aeruginosa-*induced inflammation by CF-pHBEC

To determine if MSC EVs reduce the *P. aeruginosa-*induced inflammatory response by CF-pHBEC, *P. aeruginosa* was added to the apical surface of CF-pHBECs, followed by the addition of EVs 1h later. After an additional 5h incubation, the apical and basolateral supernatants were collected for cytokine analysis. Let-7b-5p loaded MSC EVs significantly reduced *P. aeruginosa*-stimulated IL-8 secretion by 21% (p-value = 0.014), while NC MSC EVs showed a tendency to reduce IL-8 **(Figure 4A)**. However, there was no effect on IL-6 secretion by any of the treatments **(Figure 4B)**. Notably, the variation in response to treatments among donors is common when assessing drug response in pwCF^73^. However, the donors behaved similarly for the two cytokines (i.e., a donor with higher IL-8 levels also had higher IL-6 levels). qRT-PCR experiments were performed using UBC as a housekeeping/reference gene (which was unchanged by treatments (ΔCT < 1)), and mRNA fold changes were normalized to control/untreated cells. The results demonstrated that IL-8 and IL-6 mRNA were not significantly altered in the PC or MSC EV groups, as compared to the control exposed *P. aeruginosa*. This observation indicates that let-7b-5p exerts its anti-inflammatory effects by blocking translation, a typical mechanism of miRNAs^74^, or targets other genes within the IL-8 pathway **(Figures 4C - 4D)**.

**Figure 4:**
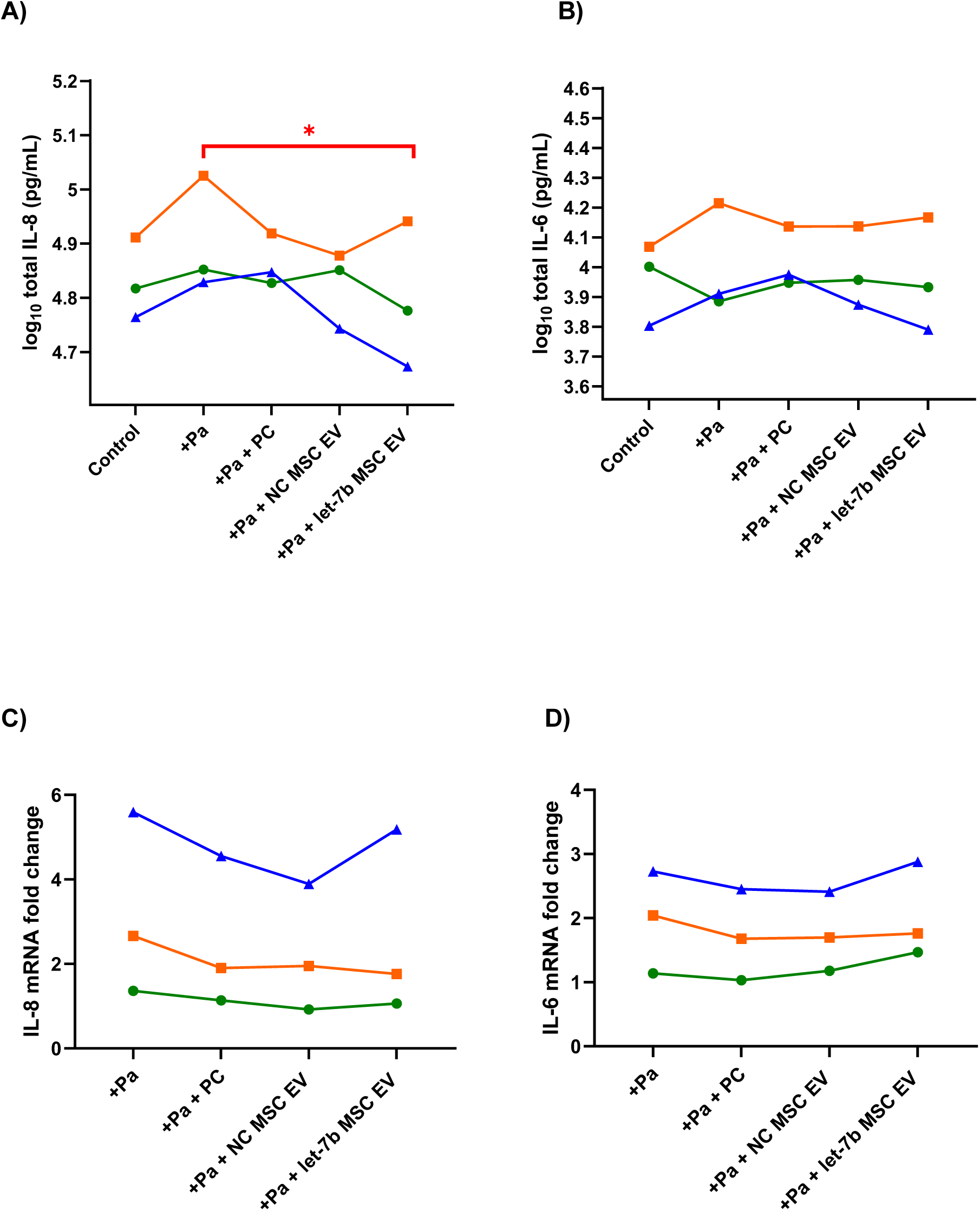
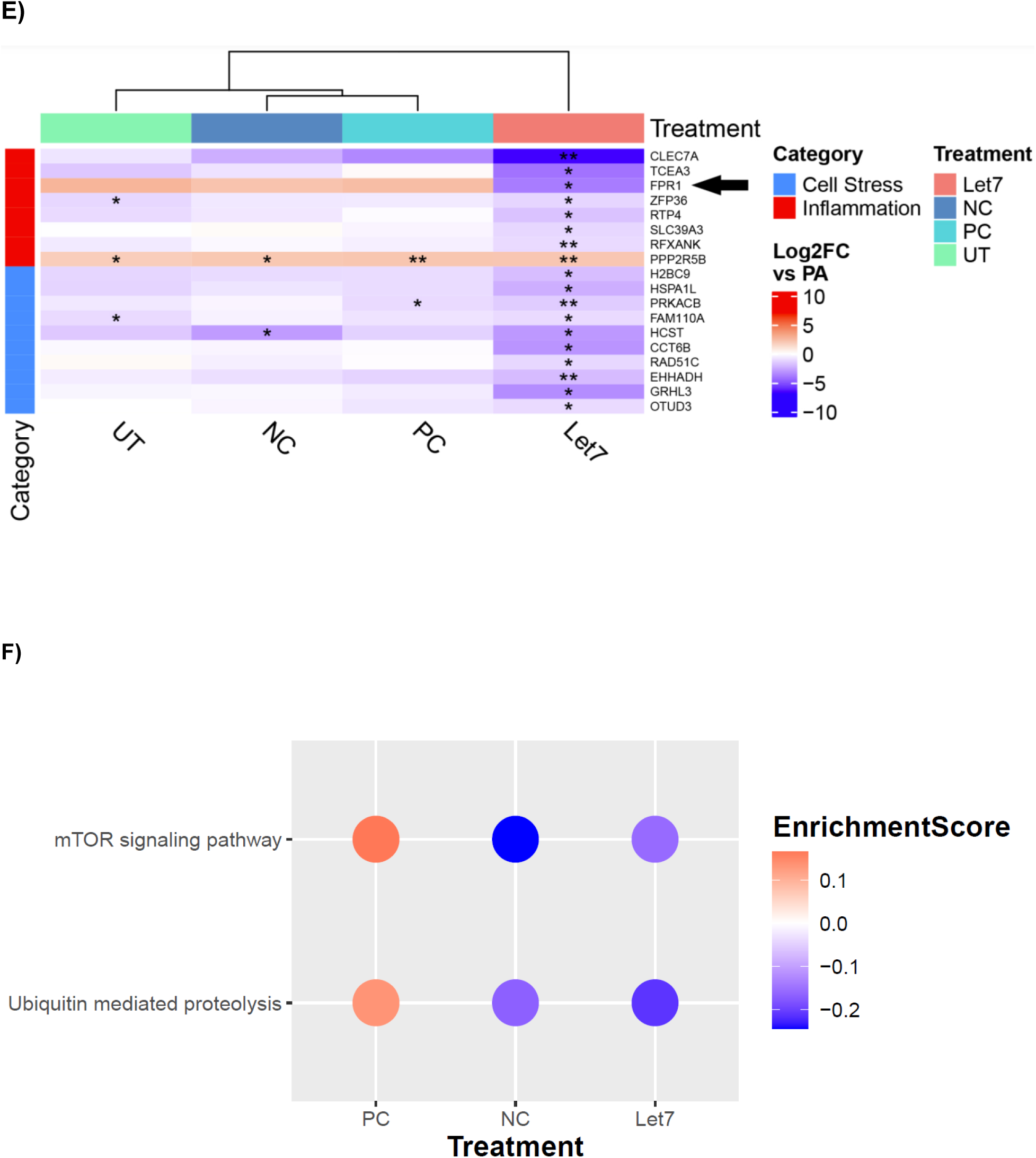
let-7b-5p MSC EVs significantly reduce *P. aeruginosa* stimulated: (A) IL-8 (p-value = 0.014) but not (B) IL-6 cytokine levels in CF-pHBEC at 6 hours. The fold changes of mRNA transcript levels of IL-8 (C) and IL-6 (D) are not significantly altered by the PC or MSC EV treatments. The 2^-ΔΔCt^ method was used to analyze relative gene expression, with UBC serving as the reference gene (the UBC ΔCT was unchanged by experimental treatment) and untreated cells as the control. Each colored line represents one CF donor across all treatment conditions. Linear mixed-effects models with donor as a random effect were used to test for statistical significance. (E) Heatmap showing significant cell stress and inflammation DEGs (differentially expressed genes with (*p* < 0.05, | log_2_FC | > 1) between let-7b-5p MSC EV treatment and *P. aeruginosa* treatment, and how they change in other conditions (untreated control, PC, NC MSC EV) relative to *P. aeruginosa* only. Red and blue represent the upregulation or downregulation of a gene, respectively, while white represents no change. *FPR1*, a predicted target of let-7b-5p is indicated with an arrow. (F) Heatmap showing the enrichment scores of the two KEGG pathways in PC, NC MSC EV and let-7b-5p exposures, all compared to *P. aeruginosa*. Red and blue represent upregulation or downregulation of a pathway, respectively. n = 3 CF-pHBEC donors (ΔF508/ΔF508); *p < 0.05, **p < 0.01; PC (process control) = unconditioned media passed through EV preparation process, NC = negative control.

To determine why let-7b MSC EVs reduced IL-8 secretion, but NC MSC EVs did not in *P. aeruginosa* exposed CF-pHBEC, we utilized bulk RNA-sequencing to examine the differentially expressed genes (DEGs) in CF-pHBEC treated with *P. aeruginosa* and let-7b MSC EVs to those treated with *P. aeruginosa* alone. Using a threshold of | log_2_FC | > 1 and p < 0.05, we identified 225 DEGs. Out of those 225 genes, we focused on inflammation/cell stress genes. Since treatments such as NC MSC EVs did not change IL-8 secretion by CF-pHBEC, we compared how these inflammation associated DEGs changed in the other treatment conditions, compared to *P. aeruginosa* alone **(Figure 4E)**. Two important observations emerged in the let-7b-5p group: (i) all but one inflammation gene and all cell stress-related genes were downregulated in the let-7b-5p MSC EV treated cells, indicating the strong therapeutic potential of let-7b-5p for pwCF and (ii) *FPR1* was the only gene significantly downregulated in the let-7b group but upregulated in the other conditions. *FPR1* encodes a receptor whose primary ligands are bacterial formylated peptides^75^ and activation of *FPR1* increases IL-8 secretion^76^ and stimulates neutrophil chemotaxis in the damaged lung^75^. Moreover, blocking *FPR1* inhibits IL-8 secretion^77^ and other inflammasome pathways^78^. Therefore, it is likely that let-7b-5p reduced IL-8 secretion by downregulating *FPR1* in CF-pHBEC. We could not find any study that linked *FPR1* downregulation with IL-6 secretion. Thus, the reduction of *FPR1* in let-7b-5p exposed CF-pHBEC, but not in NC exposed CF-pHBEC, is potentially the reason why let-7b-5b decreased IL-8 secretion. To computationally evaluate whether let-7b-5p is predicted to target *FPR1*, we employed IntaRNA^67,68^ to assess the predicted interaction energy between let-7b-5p and *FPR1*, as well as interaction energies between all other human miRNAs (2644 miRNAs) and *FPR1*. Among all miRNA candidates, let-7b-5p and *FRP1* interaction energy score was in the top 10% of all predictions (−16.48 kcal/mol), which falls well below the functional interaction threshold of -11 kcal/mol^79^ and approaches the cutoff of -16.98 kcal/mol associated with high-confidence interactions^29^. Although 209 miRNAs showed stronger predicted interaction energies with *FPR1* compared to let-7b-5p, none of them were detected in our let-7b-5p MSC EVs, or were present at extremely low levels (less than 0.001% of the miRNAs in MSC EVs)^51^. In our previous study, we had assessed the miRNA content of let-7b-5p MSC EVs, and found that let-7b-5p accounted for 97% of all miRNAs, with the remaining 3% distributed among the next 10 most abundant miRNAs^51^. None of the next top 10 miRNAs overlapped with the 209 candidates, supporting the conclusion that the observed reduction in *FPR1* expression by let-7b-5p MSC EVs was mediated by the presence of let-7b-5p.

Next, we identified signaling pathways that were differently enriched (upregulated or downregulated) in let-7b-5p exposed CF-pHBEC compared to *P. aeruginosa* alone, utilizing Gene Set Enrichment Analysis (GSEA). When comparing the enrichment scores of PC and NC MSC EV treatments (to *P. aeruginosa* alone) only two pathways were found to be enriched in opposite directions between the PC and the MSC EV groups **(Figure 4F)**. mTOR signaling and ubiquitin-mediated proteolysis were both downregulated in the let-7b-5p MSC EV and NC MSC EV groups but upregulated in the PC group. This is consistent with the pro-resolving nature of MSC EVs, suggesting that inhibition of these pathways is associated with decreased inflammation. Previous studies have demonstrated that mTOR inhibitors reduce inflammation in cystic fibrosis (CF)^80–82^, while reduced ubiquitin-mediated proteolysis has been linked to decreased inflammation and increased CFTR cell surface expression^83,84^.

### MSC EVs did not enhance the ability of CF HBEC to kill *Pseudomonas aeruginosa*

We assessed whether MSC EVs lowered bacterial burden on CF-pHBEC, potentially by increasing the intrinsic bacterial killing capacity of CF-pHBECs. Thus, 3×10^8^ CFU/mL of *P. aeruginosa* was added to the apical side of CF-pHBEC, MSC EVs were added 1 hour later, and CFU counts were measured after an additional 5 hours. Compared to the inoculum, all CF-pHBECs, irrespective of treatment conditions reduced *P. aeruginosa* CFUs from 3×10^8^ to less than 10^5^ CFU/mL in 6 hours **(Figure 5)** indicating that CF-pHBEC had an intrinsic ability to kill *P. aeruginosa* over time. However, MSC EVs had no effect on the bactericidal effect of CF-pHBEC on *P. aeruginosa* after 6 hours of exposure **(Figure 5)**.

**Figure 5:**
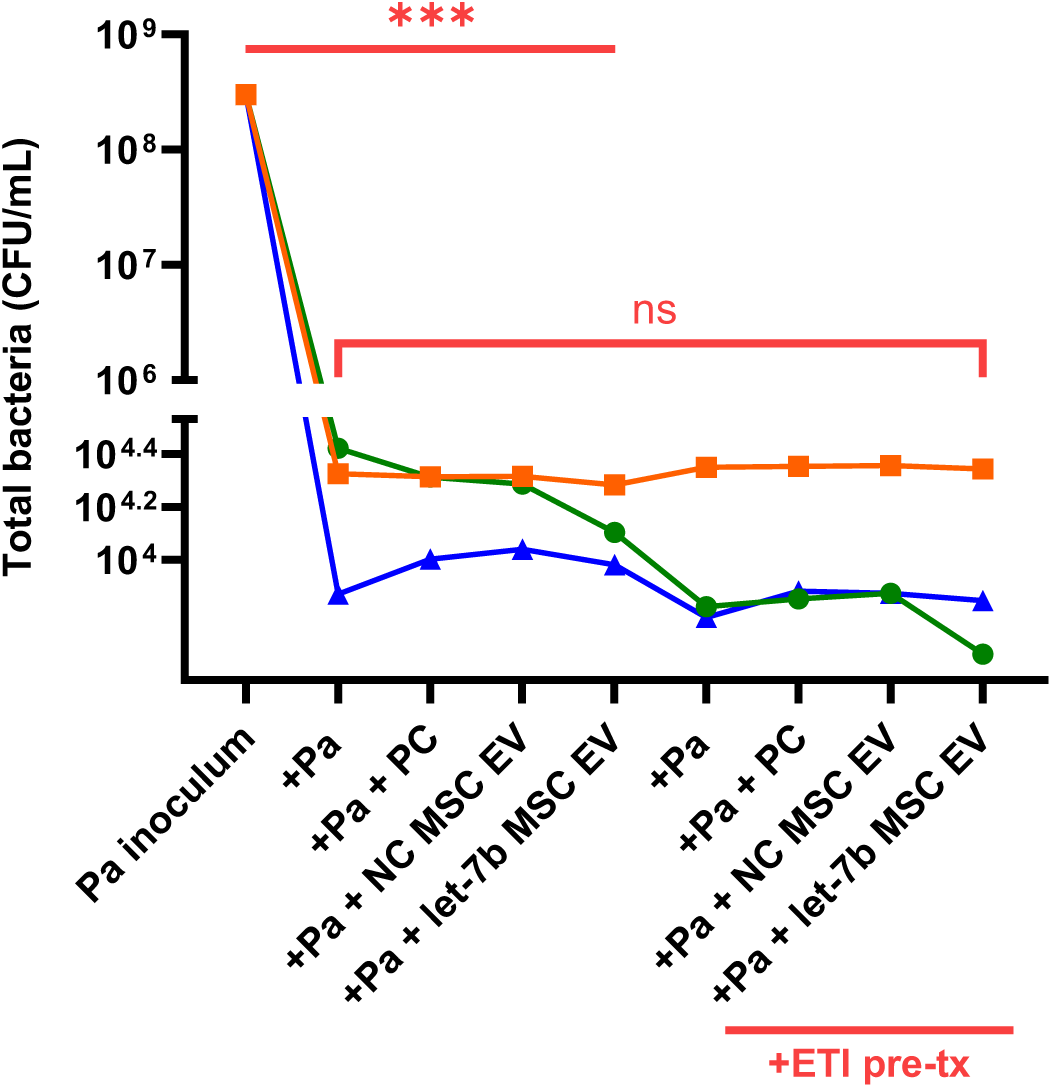
CF-pHBEC from three donors kill *P. aeruginosa* (Pa) after 6 hours. The CFUs fell from 3 × 10^8^ CFU/mL Inoculum) to < 4.4×10^4^ CFU/mL. Each colored line represents one CF donor across all treatment conditions. Linear mixed-effects models with donor as random effect were used to test for statistical significance in the bacterial burden assay; n = 3 CF-pHBEC donors (ΔF508ΔF508); ***p < 0.001 comparing *P. aeruginosa* (Pa) inoculum to all four treated CF-pHBEC groups ± ETI; PC (process control), NC = negative control.

### ETI enhances the bactericidal effect of some CF-pHBEC

Given that MSC EVs did not enhance bacterial killing by CF-pHBEC, we tested the hypothesis that ETI may enhance the bactericidal effect of CF-pHBEC on *P. aeruginosa*. ETI boosted the intrinsic bactericidal capacity of CF-pHBECs in some donors **(Figure 5).** For example, bacterial killing was pronounced in one donor (green line), mild in another other (blue line), and absent in the remaining donor (orange line). These variations indicate donor-specific responses to ETI, which has been reported in pwCF^54,85^. In addition, in the presence of ETI, neither PC nor NC MSC EVs nor let-7b-5p MSC EVs had a significant effect on *P. aeruginosa* compared to PC.

## DISCUSSION

In this study, we demonstrate that all MSC EVs, including let-7b-5p-MSC EVs, completely inhibited *Pseudomonas aeruginosa* biofilm formation on pHBECs. This observation indicates that MSC EVs are likely to be a highly beneficial therapeutic that overcomes antibiotic-resistant biofilms^6,7^. Additionally, let-7b MSC EVs decreased *P. aeruginosa*-induced expression of several proinflammatory genes. Further, let-7b MSC EVs reduced IL-8 secretion by CF-pHBECs potentially in part by reducing *FPR1* expression. Previous studies have shown that *FRP1* regulates IL-8 secretion^76,77^. MSC EVs (NC and let-7b-5p MSC EVs) also reduced the mTOR and ubiquitin mediated proteolysis pathways. Taken together, we conclude that MSC EVs in general and MSC EVs engineered to contain let-7b-5p will have potent anti-inflammatory effects in pwCF infected with *P. aeruginosa,* which is predicted to reduce the damaging effects of lung inflammation in CF. Moreover, in some CF-pHBECs, ETI increased the bactericidal activity of CF-pHBEC towards *P. aeruginosa*. These results are consistent with our recent study in CF mice in which MSC EVs reduced inflammation and decreased the bacterial abundance in the CF murine lung^51^, indicating that the effect of MSC EVs in mice is due in part to effects on bronchial epithelial cells

Our earlier research revealed a reduction in bacterial burden and immune cell infiltration in the lungs of CF mice treated with MSC EVs and let-7b-loaded MSC EVs^51^. These findings in mice suggest multiple mechanisms contributing to the observed *in vivo* effects. One plausible mechanism is the direct bactericidal activity of MSC EVs, as established in previous studies^86,87^. The current study provides evidence for an additional mechanism: MSC EVs eliminate *P. aeruginosa* biofilm formation on pHBEC, thereby enhancing the host’s ability to clear infections in response to antibiotics, since biofilm bacteria are highly resistant to antibiotics^88^. Inhibition of biofilm formation by MSC EVs is particularly significant, as biofilms are a key factor in the persistence of antibiotic-resistant *P. aeruginosa* in the CF lung. Our study is in general agreement with reports demonstrating that conditioned media from MSC inhibits the formation of abiotic *S. aureus* and *E. coli* biofilms^89^, and that the equine MSC secretome, which contains EVs, inhibits abiotic biofilm formation by *P. aeruginosa*, *S. aureus*, and *S. epidermidis*^90^. Importantly, our study advances these reports by demonstrating that MSC EVs reduce the formation of *P. aeruginosa* biofilms growing on pHBEC, a model that typically supports robust biofilm formation^91^. Moreover, let-7b-5p loaded MSC EVs reduced IL-8 secretion, an effect, that in part, can account for the reduction in immune cell recruitment to the lungs of CF mice exposed to MSC EVs^51^. However, MSC EVs, regardless of let-7b loading, did not enhance the intrinsic bacterial killing capacity of CF pHBECs under the conditions tested.

An important question raised by our findings is why did MSC EVs not directly lower the bacterial burden on CF-pHBEC, despite their established antibacterial properties^39,43,44^. One potential explanation is that the 6-hour time point used in our experiments may have been insufficient for the EVs to fully exert their effects. For instance, a previous study demonstrated that MSCs directly killed bacteria via the secretion of the antimicrobial peptide LL-37; however, this effect required 16 hours^92^, considerably longer than the 6 hour exposure in the present study. Unfortunately, *P. aeruginosa* begins to have cytotoxic effects in our model system after 6 hours of exposure precluding extended time points. Additional studies, beyond the goals of this report, are required to develop a model to determine the time of exposure to MSC EVs required to reduce infection by *P. aeruginosa*. Notably, in our mouse studies MSC EVs reduced the lung burden of *P. aeruginosa* on 3 days after exposure to MSC EVs ^51^.

This study offers several significant advantages. First, considering the growing urgency to develop anti-infectives in a world of escalating antibiotic resistance, we are the first group to show that MSC EVs inhibits the biofilm-forming capacity of *P. aeruginosa* on pHBEC. Second, we compared EVs from two sources of passage-matched human bone marrow-derived MSCs and characterized them using multiple approaches, as recommended by the International Society of Extracellular Vesicles^72^. Interestingly, while phenotypically similar, EVs derived from ATCC MSCs were more effective in reducing inflammation than those from PACT MSCs. MSCs from different sources often excel in distinct functions, depending on the specific metric being assessed. For example, Bonfield et al. reported that MSC from different donors had variable effects against specific pathogens, with one donor showing greater efficacy against *Pseudomonas* and another against *Mycobacterium*^93^. Another strength of our study is the use of primary CF-pHBECs instead of cell lines, which are derived from a single donor. Moreover, the use of primary cells is more physiologically relevant, as they produce mucus, which affects bacterial virulence^94,95^. Our experiments also demonstrate the ability of MSC EVs to diffuse through the mucus overlying CF-pHBEC and exert their effects^96,97^. Additionally, we observed donor-specific variability in the response to EV treatment, underscoring the importance of personalized medicine^98^. For instance, one donor exhibited a hypo-responsive profile to MSC EVs, whereas others responded more robustly, highlighting intrinsic differences that were accounted for by modeling the donor as a random effect in our statistical analyses. Notably, pwCF, even those with the same mutation, manifest a wide range of phenotypes, infection profiles and responses to HEMT ^54,73,85^.

Despite these advances, our study has a few limitations that can be addressed in the future. While we examined EVs from two MSC sources, expanding the donor pool could identify donors whose EVs are superior to the ones from ATCC. This would be particularly beneficial for clinical translation and a personalized medicine approach. Alternatively, one can also test the efficacy of EVs obtained from iPSC-derived MSCs. Additionally, we used a fixed dose of MSC EVs, selected based on prior measurements in airway epithelial cell culture supernatants and human bronchoalveolar lavage fluid^56,57^. Future studies exploring the dose and time dependent effects of MSC EVs would be important to develop MSC EVs as a therapeutic. In addition, RNA sequencing revealed that let-7b-5p is responsible, at least in part, for the MSC EV induced decrease in *FPR1* gene expression and potentially the associated reduction in inflammation. However, we cannot rule out the possibility that other miRNAs or constituents of MSC EVs play a role in the antibacterial and anti-inflammatory effects observed. Finally, although others have shown a direct relationship between *FRP1* and IL-8 secretion, additional studies, beyond the goals of this study, are required to demonstrate that in pHBEC let-7b-5p decreases FRP1 protein abundance and signaling.

In summary, we have demonstrated that MSC EVs exhibit both anti-biofilm and anti-inflammatory effects on CF-pHBEC, and that let-7b-5p has several unique anti-inflammatory effects. These findings suggest that engineered MSC EVs represent a promising therapeutic candidate to reduce both infection and inflammation in pwCF, even in those treated with HEMT. By leveraging bioinformatics, our Rocket-miR application (a program that predicts that let-7b-5p and other miRNAs that have antibacterial effects on 24 pathogens, including all ESKAPE pathogens ^29^) and synthetic engineering, our long-term goal is to develop a dual-function anti-infective and anti-inflammatory approach to eliminate chronic infection of all pathogenic bacteria and inflammation in pwCF.

## ACKNOWLEDGEMENTS

This study was supported by the Cystic Fibrosis Foundation (STANTO19G0, STANTO20P0, STANTO23R0, and STANTO19R0) and the National Institutes of Health (P30-DK117469 and R01HL151385) to BAS. We thank RoosterBio for analysis of the MSC EVs (Table 1). We thank Dr. Carey Nadell (Dartmouth College) for the PA14-mK02. Cells and media were provided by the Marsico Lung Institute Tissue Procurement and Cell Culture Core supported by NIH grant DK065988 and CF Foundation grant BOUCHE19R0. RNA-seq was carried out in the Genomics and Molecular Biology Shared Resource (RRID:SCR_021293) at Dartmouth which is supported by NCI Cancer Center Support Grant 5P30CA023108 and NIH S10 (1S10OD030242) awards. Support for RNA-seq data analysis was provided through Dartmouth Cancer Center Bioinformatics & Biostatistics Shared Resource (BBSR) supported by NCI award P30CA023108 and Dartmouth’s Center for Quantitative Biology (CQB) Genomics Data Sciences Core (GDSC) supported by NIH award 3P20GM130454, and NIH F31HL172440 to LAC.

## DATA AVAILABILITY

Data will be made available upon reasonable request. RNA sequencing data has been uploaded to NCBI GEO (Accession number: GSE295607).

## DISCLAIMERS

The funders had no role in study design, data collection and analysis, publication decisions, or manuscript preparation.

## AUTHOR CONTRIBUTIONS

S.S., R.B., A.N., M.J.W., D.J. W., T.L.B., and B.A.S. conceived and designed research; S.S., R.B., and A.N. performed experiments; S.S., L.A.C., and L.T. analyzed data; S.S., R.B., A.N., L.A.C., L.T., and B.A.S. interpreted results of experiments; S.S. and L.A.C. prepared figures; S.S. and B.A.S. drafted the manuscript; all authors edited and revised the manuscript; all authors approved the final version of the manuscript.

